# Resource allocation strategies and mechanical constraints drive the diversification of stick and leaf insect eggs

**DOI:** 10.1101/2023.11.10.566544

**Authors:** Romain P. Boisseau, H. Arthur Woods

## Abstract

The diversity of insect eggs is astounding but still largely unexplained. Here, we apply phylogenetic analyses to over 210 species of stick and leaf insects (order Phasmatodea), coupled with physiological measurements of metabolic rate and water loss, to evaluate several major classes of factors that may drive egg morphological diversification: life history constraints, material costs and mechanical constraints, and ecological circumstances. We show support for all three classes, but egg size is primarily influenced by female body size and strongly trades off with egg number. Consequently, females that lay relatively fewer but larger eggs, which develop more slowly because of disproportionately low metabolic rates, tend to bury or glue them in specific locations, instead of simply dropping them from the foliage (ancestral state). This form of parental care then directly favors relatively elongated eggs, which may facilitate their specific placement and allow easier passage through the oviducts in slender species. In addition, flightless females display a higher reproductive output and consequently lay relatively more and larger eggs compared to flight-capable females. Surprisingly, local climatic conditions had only weak effects on egg traits. Overall, our results suggest that morphological diversification of stick insect eggs is driven by a complex web of causal relationships among traits, with dominant effects of resource allocation strategies and mechanical constraints.

## Introduction

Insect eggs share a common set of characteristics defined by their function: they are propagules -- finite packages of resources and information that support embryos as they grow from single cells into complex individuals that hatch into free-living juveniles^1^. Across taxa, eggs are recognizable as such because they share deep homologies. Egg diversity, by contrast, is more difficult to explain. Eggs diverge enormously in size, shape, composition, structure, physiology, and duration of development ^2^. What rules structure this diversity? How universal or taxon-specific are they? These questions have a long history of study in other taxa, especially birds ^3^, reptiles ^4^, fish ^5^, and marine invertebrates ^6,7^, though most studies have focused just on egg size. Below, we briefly outline three non-exclusive classes of hypotheses on the major causes of egg diversification in insects ^8^.

### Size, life history, and pace of life

An organism’s size can predict many other aspects of its life history, including length of development, potential fecundity, metabolic rate, and adult lifespan. In eggs from insect and non-insect taxa, for example, larger adults typically lay larger eggs ^5,9,10^, and large eggs typically develop more slowly than small eggs ^9,11–16;^ but for an alternative finding on insects, see ^8^. In addition, like scaling relationships in other taxa and other life stages ^17,18^, metabolic rates of eggs scale hypoallometrically with egg mass ^19^. Larger adults thus tend to lay larger eggs that develop over longer periods of time, supported by disproportionately low metabolic rates. In addition, egg size may trade off with egg number; females have access to finite resources, and even if they acquire nutrients in adulthood, they face the proximate problem of whether to construct smaller numbers of larger eggs or greater numbers of smaller eggs. Egg size- number tradeoffs in insects have been demonstrated both intraspecifically ^20–22^ and interspecifically^10,23,24^.

### Mechanical constraints on egg size and shape

Eggs may face strong geometric constraints on shape as a function of size ^3,8^. For example, because eggshell materials are costly (typically rich in protein), females laying more or larger eggs may minimize costs by producing more spherical eggs ^4,10^. On the other hand, larger eggs may need to have higher aspect ratios so that they can pass more easily through the reproductive canal during oviposition ^4,10^. Larger eggs may also require higher ratios of surface area to volume, allowing them to obtain oxygen at rates high enough to support embryonic metabolism and to minimize diffusion distances between surface and central tissues ^25^. In addition, supporting higher metabolic rates may require higher-conductance eggshells, which in turn can lead to higher rates of water loss ^16,26^.

### Response to ecological circumstances

Egg traits may evolve in response to local suites of ecological conditions that they experience, including: (1) patterns of predation in relation to size, placement, or camouflage of the eggs, or as functions of the mechanical and chemical defenses that they deploy ^27,28^; (2) whether females are flight-capable or flightless ^3^; and (3) local environmental conditions in their microsites, including whether eggs are aquatic or aerial ^8^.

Church, Donoughe et al. ^8^ recently examined a subset of these possibilities using data on egg and adult traits derived from literature on over 6,700 species in 526 families distributed across all extant orders, although not all traits were available for all species. Using phylogenetic analyses, they reached three main conclusions. First, geometric constraints on egg diversification were detectable but weak; larger eggs, for example, tended to have higher aspect ratios. Second, larger females indeed tended to lay larger eggs, but, contrary to other studies, those eggs did not have systematically longer developmental periods. Third, the most important drivers of changes in egg size and shape were the ecological circumstances of oviposition. In particular, evolutionary shifts from aerial oviposition (onto surfaces exposed to the atmosphere) to aqueous oviposition (into water or body fluids) led to systematic reductions in egg size and changes in shape.

Because of the large data set used, the analyses of Church, Donoughe et al. ^8^ are uniquely comprehensive and powerful. Nevertheless, some conclusions – e.g., that larger eggs do not take longer to develop – are surprising and may reflect that some analyses were constrained by lack of data on focal traits to a relatively small number of species distributed across multiple major clades of insects. Such a situation may increase the probability that patterns in focal traits, even if present, are obscured by other evolved differences among major clades (i.e., orders). A complementary approach would be to focus on individual clades with extensive sampling of species from across the group phylogeny. Here we present such an analysis for stick and leaf insects (order Phasmatodea), which appear to have originated in the early Cretaceous and diversified extensively following the K-T boundary ^29,30^ (but see ^31^). Worldwide there are about 3,400 described species^32^, for which we have compiled data on morphological and developmental traits of eggs and adults on nearly 210 species from approximately 27% of the ∼520 described genera.

As masters of crypsis and masquerade, stick and leaf insects show remarkably diverse body morphologies, ranging from elongated stick-like silhouettes to robust, stocky form^33^. They also produce remarkably diverse eggs, which span wide ranges in size, shape and structure. Eggs of some Heteropteryginae, for example, are among the largest produced by any insect, reaching 300 mg and 12 mm in length in *Haaniella echinata*^34^. By contrast, eggs of *Spinoparapachymorpha spinosa* (Clitumninae) are only 2 mg and 1.6 mm in length^35^.

In Euphasmatodea, the evolution of a hardened egg chorion is a key innovation and may have contributed to the group’s extreme diversification by opening up multiple routes of unusual dispersal ^30,36^. Hard shells and their associated structures can withstand hazardous falls from the forest canopy, float for extended periods of time on sea water (hydrochory)^37^, bear ant-attractive capitula (analogous to plants’ eliasomes, myrmecochory)^38^, and even survive passage through the guts of birds (endozoochory) ^39,40^. Accordingly, female phasmids employ a variety of egg-laying strategies, including passively dropping or actively flicking eggs to the forest floor, burying them in soil or other soft substrates, and gluing them to plant surfaces individually, in groups, or inside complex ootheca^30,41^. Since the late 1800s^42–44^, biologists have suggested that the morphological resemblance of phasmid eggs to seeds is a form of mimicry or masquerade. While this resemblance is indeed impressive, its ecological and evolutionary significance is still largely unknown.

Owing to their extreme diversity, relatively large size, and ease of breeding in captivity, stick insects are commonly kept as pets by amateur breeders or in classrooms, which has facilitated the compilation of a large data set on egg size, shape and physiology^35,45,46^. We leverage this comparative data set in a phylogenetic context, along with additional data on rates of metabolism and water loss by eggs of five of the species, to evaluate the relative importance of the diversifying factors proposed above. Overall, signatures of many factors appeared in patterns of egg diversification. Nevertheless, causal relationships (derived from phylogenetic path analyses) between these factors revealed that egg size was mostly affected by resource allocation strategies and trade-offs, while egg shape was mostly influenced by mechanical constraints.

## Methods

### Phylogenetic reconstruction

For the present study, we used a phylogeny from a companion study dealing with morphological convergence in adult female stick insects^33^, which includes a total of 314 phasmid taxa (9% of phasmid species diversity and 33% of currently recognized generic diversity) and one embiopteran species as outgroup (i.e., from the sister clade of Phasmatodea). The phylogenetic analyses were originally performed using genetic data from 3 nuclear (*i.e.,* 18S rRNA, 28S rRNA and histone subunit 3) and 4 mitochondrial genes (*i.e.,* 12S rRNA, 16S rRNA, cytochrome-c oxidase subunit I and II) and Bayesian inferences (BEAST2, v. 2.6.3,^47^. The basal backbone topology of the tree was constrained to match that of transcriptome-based studies that could confidently infer the deep relationships between all the major clades of Phasmatodea ^29,31^. The resulting maximum clade credibility tree was overall strongly supported and congruent with previous phylogenetic reconstructions^30,33,48,49^.

### Egg morphology

We collected high quality photographs of eggs in dorsal and lateral view for a total of 144 different species included in the phylogeny mainly from the egg picture database of F. Tetaert (Office pour les insectes et leur environnement OPIE, France, retrieved from phasmatodea.fr in August 2021) and from the published literature and other online databases (see Dataset S1 for details). We applied the guided landmark-based methodology of Church et al. ^50^ to quantify egg shape traits (Fig. S1A). We used the R software (v4.1.1)^51^ and the package “raster” ^52^, to measure egg length (L) from the base of the operculum to the posterior end of the egg, and width and height along three different latitudinal lines at ¼, ½ and ¾ (respectively w_1_, w_2_, w_3_ and h_1_, h_2_, h_3_) of the egg longitudinal axis (Fig. S1A). Egg width (w) and height (h) were considered as the maximum values of w_1_, w_2_, w_3_ and h_1_, h_2_, h_3_ respectively. Egg volume was then estimated using the equation for the volume of an ellipsoid 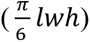. We verified the relevance of using this estimate of volume as our measure of egg size by regressing egg volume on egg mass (after log_10_- transformation) for species where that information could be collected (n = 85 species, see Dataset S1 for details). We found a strong correlation between the two (*β*=0.87 ± 0.04, p<0.0001, R^2^=0.84, Fig. S2) and therefore chose to use egg volume as our measure of egg size as it was available for more taxa (n=144 species). Egg surface area (SA) was calculated as the surface area of an ellipsoid using the approximate formula:

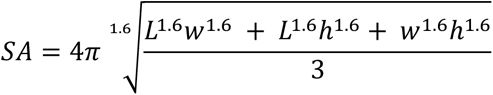

To characterize egg shape variation while controlling for size, we performed a phylogenetic Principal Component Analysis (pPCA, *R function*: “R package”; *phyl.pca*: “phytools” ^53^) including L, w_1_, w_2_, w_3_, h_1_, h_2_ and h_3_, but original values were substituted with residuals calculated from a phylogenetically-corrected generalized least-squares (PGLS) regression against egg volume (*gls*: “nlme” and “ape” ^54,55^, lambda=’ML’).

### Adult female morphology

To quantify adult female size, we used data on adult female volume originally collected in a previous study (n = 212 species)^33^, which estimated female body volume as an ellipsoid using the average body length, width, and height of a species. Original measurements were obtained from digital images of live and dried specimens. From this study, we also used the species average width of the female’s ninth tergite (i.e., width the ninth abdominal dorsal plate) as it roughly corresponds to the location of the oviduct, from which eggs emerge during oviposition ^56^ and whose diameter may mechanically constrain egg morphology.

### Ecological variables

Using primarily the dataset assembled by Robertson et al. ^30^, we classified egg oviposition strategies into three categories: females drop or flick eggs to the ground from higher up in the local canopy, bury eggs into soil or other soft substrates, or glue eggs to substrate (including eggs encased in an ootheca). We then mapped oviposition strategies onto the phylogeny and ran an ancestral state reconstruction using stochastic character mapping (*make.simmap*: “phytools”). We calculated the transition matrix using MCMC and assumed different transition between all states (model “ARD”). We subsequently simulated and summarized 1,000 stochastic character maps to obtain posterior probabilities for each state at each node.

We gathered observational data on the flight capacities of adult females across species. Female flight capacity was classified as either flight-capable (including gliding) or flightless (including parachuting)(Dataset S1, n = 215 species).

We also collected climate data for each species (n = 214) based on current geographic distribution and corresponding to climate conditions experienced at the most central location of the range where the species have been collected (using the collection location of type specimens) or observed (using observations on iNaturalist, available from https://www.inaturalist.org, accessed July 2021). The dataset included annual mean temperature, mean diurnal range (i.e., mean of monthly (maximum - minimum temperature)), temperature seasonality (i.e., standard deviation × 100), annual temperature range (i.e., maximum temperature of warmest month - minimum temperature of coldest month), annual precipitation, precipitation seasonality (i.e., coefficient of variation) from worldclim ^57^. The dataset also comprised the length of the growing period (number of days during a year when temperatures are above 5°C and precipitation exceeds half the potential evapotranspiration, available from https://data.apps.fao.org/map/catalog^58^), net primary production of biomass (grams of dry matter per m^2^ per year; Climate Research Unit, Univ. of East Anglia, period 1976-2000, FAO Map Catalog, available from https://data.apps.fao.org/map/catalog), and total annual growing degree days (a measure of the annual amount of thermal energy available for plant and insect growth; Climate Research Unit, Univ. of East Anglia, available from https://sage.nelson.wisc.edu/data-and-models). We then ran a principal component analysis to summarize climatic variation across the geographic distribution of the phasmids (*prcomp*: “stats”). We kept the first two axes of the PCA (climate PC1 and PC2, respectively, explaining 56% and 17% of the total variation) to quantify climatic variation between species (Fig. S3). Climate PC1 reflected tropical versus temperate patterns differing in annual precipitation, annual temperature, and magnitude of diurnal and annual variation in temperature and consequently in net primary productivity (i.e., food availability for herbivorous insects). PC1 is overall high in tropical regions and low in more temperate and seasonal regions with restricted growing periods. Climate PC2 mostly represented variation in precipitation seasonality with regions showing strong shifts between dry and wet/monsoon seasons displaying a high PC2.

### Life history variables

We collected data on the average number of eggs laid by a female during its lifetime (n = 100 species) and on average embryonic development duration (from oviposition to hatching, n = 136 species) from the published literature, amateur breeding guides and the phasmid breeder community (Dataset S1). However, it should be noted that information on egg incubation time is often quite approximate (±2 weeks) and does not account for temperature. However, phasmids are typically bred at room temperature (20-24°C) and likely show higher variation in development time across species (from 1.25 to 11.5 months) relative to measurement error. Lifetime reproductive output was calculated as the product of egg volume and the fecundity (n = 96 species with sufficient data) and was used to quantify average female lifetime reproductive investment.

### Phylogenetic analyses and hypothesis testing

We tested hypotheses on the drivers of egg diversification in phasmids following the framework outlined in the introduction and organizing the hypotheses into three classes. All analyses were performed in R (v4.1.1)^51^. Continuous variables were log_10_-transformed prior to the statistical analyses. PGLS models were run using the R packages “nlme” and “ape” (correlation= “corPagel”)^54,55^. Pagel’s lambda was estimated using maximum likelihood. Significance of the effect of each explanatory variable was evaluated using a type I (sequential) analysis of variance (ANOVA) or analysis of covariance (ANCOVA). The significance of interaction terms was systematically tested and non-significant interaction terms (p<0.05) were removed from the final models to improve the estimation of intercepts and effect sizes. When appropriate, post hoc pairwise comparisons were run to contrast the intercepts and/or slopes of different levels of a categorical explanatory variable (*emmeans* and *emtrends* : “emmeans”)^59^ using the Holm method to account for multiple testing.

First, we investigated hypotheses related to the effect of life history strategies on egg size. We tested whether egg size was dependent on adult female size, whether egg size traded off with egg number, and whether females varying in oviposition mode (i.e., parental care investment) varied in reproductive allocation strategy. We examined the scaling relationships between adult female size and egg volume and fecundity by running PGLS regressions including lifetime fecundity or egg volume as the response variables and female body volume and oviposition mode as the explanatory variables. Then, to specifically test for a trade-off between egg number and size, we ran a PGLS model including fecundity as the response variable and female body and egg volume as explanatory variables, predicting a negative effect of egg volume on fecundity after accounting for female body size. We then compared the total reproductive output of females as a function of female body volume and oviposition mode, predicting that females investing more energy in parental care (by burying or gluing eggs to specific plants or substrates) would have relatively lower reproductive output. Finally, we tested whether larger eggs developed more slowly than smaller eggs by building a PGLS model including duration of embryogenesis as the response variable and egg volume and oviposition mode as predictors.

Next, we tested hypotheses related to mechanical and geometric constraints on egg size and shape. When egg size increases, eggs may become wider to save on costly eggshell material, leading to a hyperallometric relationship between egg width and length (i.e., slope greater than 1 on a log-log scale). Alternatively, as they get larger, eggs may become more elongated to be able to fit through a narrow opening during oviposition, to increase their surface area to volume ratio to obtain relatively more oxygen, and/or to minimize diffusion distances between the surface and the central tissues. In this case, we expect a hypoallometric relationship between egg width and egg length (i.e., slope lower than 1 on a log-log scale). We examined the scaling relationship between egg width and egg length using a PGLS regression including egg width as the response variable and egg length and oviposition mode as explanatory variables. Slopes were compared to isometry using 95% confidence intervals. We also tested the scaling relationship of egg surface area and egg size by including egg surface area as the response variable and egg volume and oviposition mode as explanatory variables in a PGLS model. Finally, we examined the relationship between egg width and the width of the female’s ninth tergite (where the opening of the oviduct is located) by running a PGLS model including egg width as the response variable and female ninth tergite width and oviposition mode as the explanatory variables. To investigate whether females with relatively narrower abdomen tips laid relatively more elongated eggs, we obtained measures of egg width relative to egg length, and of female ninth tergite width relative to female size by extracting the residuals of two PGLS regressions. One had egg width as a function of egg length and the other had ninth tergite width and female volume. Then we ran a PGLS model including residual egg width as a function of residual ninth tergite width and oviposition mode.

Finally, we examined hypotheses related to ecological circumstances. Oviposition mode is expected to affect egg size and shape as the conditions experienced by dropped, buried or glued eggs are likely to be very different in terms of exposure to predators, oxygen availability, and desiccation risk. Buried eggs may benefit from being more elongated so that females can insert them more easily into the substrate and so that they have relatively more surface area for gas exchange underground. Glued eggs, by contrast, are expected to have shapes that vary depending on the substrate to which they are glued (e.g., thin grass leaf, bark, broad leaf). Additionally, glued eggs are likely to be more exposed to predators and desiccation than eggs on or into soil and may therefore be selected to develop more rapidly. We also tested whether flight capacity affected egg size evolution, taking advantage of the numerous transitions between flight-capable and flightless species seen in stick insects ^48,60–62^. Because flight is costly ^63^, we predicted that the reproductive investment by flying females would be relatively lower. We also looked at the effect of climate on egg size and shape. We hypothesized that drier climates would drive the evolution of rounder, larger eggs to increase total water stores and limit surface area to volume ratio and therefore relative rates of water loss. In order to test the effect of these ecological variables on egg size, we built a PGLS model including egg volume as the response variable and female body volume, fecundity, oviposition mode, female flight capacity, climate PC1 and PC2 as predictors. Thus, variation in fecundity and female size was accounted for to avoid confounding effects. Then we built similar PGLS models with either egg surface area, egg width, reproductive output and duration of embryogenesis as response variables and respectively egg volume, egg length, female body volume and egg volume as the first predictors to account for size effects. All models then included oviposition mode, female flight capacity, climate PC1 and 2 as predictors.

### Phylogenetic path analyses

We used phylogenetic confirmatory path analyses to compare fits of prespecified models of causal hypotheses among variables to our data, while accounting for nonindependence of observations due to phylogenetic relatedness among species (R package “phylopath”) ^64–66^. The variables included in the models were egg volume (n=142 species), egg shape PC1 (i.e., elongation; n=142), female lifetime fecundity (i.e., egg number; n=95), adult female body volume (n=142), embryonic development time (n=120), female relative ninth abdominal segment width (residuals of PGLS regression between adult female ninth tergite width and body volume; n=142), climate PC1 (i.e., temperate versus tropical; n=142), female flight capacity (binary: 1= flight capable, 0= flightless; n=142) and parental care (binary: 1= eggs placed in specific locations (buried or glued), 0= eggs dropped; n=142). All continuous variables (except principal components and residuals) were log_10_-transformed. Models with different configurations of these variables were compared using the C-statistic Information Criterion corrected for small sample size (CICc) and were considered equivalent if ΔCICc <2. Models were designed to test the existence and directionality of effects between the variables. We started from an initial model including many logical links (arrows) between variables (Fig. S4). From this initial model we went through a model selection procedure consisting of 3 phases. During this process, we started from a main model and either dropped specific arrows or reversed their directionality one by one. All the produced models were then compared, and the arrow change associated with the best model was applied to all models. The process was then repeated until the main model was equivalent to the best model. In the first phase, we dropped arrows one by one. In the second phase, we changed their direction and in the third phase we dropped arrows again. Each phase started from the main model produced by the previous phase. In phase 2, we excluded models that made little logical sense. For example, climate PC1 was never considered a consequence of any other variable; female size was never considered a consequence of egg shape, parental care and egg number; and egg size was never considered a consequence of embryonic development time. 3 alternative models produced cyclic graphs which are incompatible with path analyses. For these, we also switched the directionality of another arrow to reestablish acyclicity: female flight was considered a consequence of both egg size and number, parental care was considered both a consequence of both egg size and number and finally relative female abdominal width was considered a consequence of both egg shape and size (Fig. S5). At the end of phase 3, the average of the best performing models (ΔCICc < 2) was calculated, and path coefficients were averaged only over models where the path exists (avg_method=“conditional”) ^66^.

### Study animals for physiological experiments

Sixteen freshly laid eggs were obtained from five unrelated phasmid species from culture stocks: *Eurycantha calcarata* (Lucas, 1869; subfamily Lonchodinae), *Extatosoma tiaratum* (Macleay, 1826; subfamily Phasmatinae: “Lanceocercata”), *Heteropteryx dilatata* (Parkinson, 1798; subfamily Heteropteryginae), *Medauroidea extradentata* (Brunner von Wattenwyl, 1907; subfamily Clitumninae) and *Phyllium philippinicum* (Hennemann, Conle, Gottardo & Bresseel, 2009; subfamily Phyllinae). All eggs were fertilized except *M. extradentata* eggs, which were obtained from a parthenogenetic culture. *P. philippinicum*, *M. extradentata* and *E. tiaratum* females typically drop or flick their eggs away during oviposition while *E. calcarata* and *H. dilatata* bury them ^30^. Throughout embryonic development, eggs were kept individually in separate wells of a plastic 24-well plate placed inside a plastic box whose bottom was filled with water to maintain nearly 100% relative humidity around the eggs. The eggs were kept at room temperature (21 - 23 °C) under a natural day/night cycle until hatching.

### Egg metabolic rates

Egg metabolic rate was estimated as the rate of CO_2_ production using flow- through respirometry. CO_2_ concentrations were measured using a Licor LI-7000 CO_2_/H_2_O analyzer (Licor, Lincoln, NE, USA) in differential mode associated with a 16-chamber flow-through respirometry and data acquisition system (MAVEn-FT, Sable Systems International, North Las Vegas, NV, USA). The analyzer was calibrated frequently using CO_2_-free air and 25 ppm CO_2_ in N_2_ (NorLab, Norco, Boise, ID, USA). Air flow rates were set to 25 ml.min^−1^ (standard temperature and pressure) in the 16 experimental chambers. Gases circulated between the instruments in 3 mm inner-diameter plastic tubing (Bevaline-IV, Cole Parmer, Vernon Hills, IL, USA). Dry, CO_2_-free air was first directed through a column containing Drierite and Ascarite to trap residual CO_2_, then through the reference cell of the analyzer (referred to as cell A), which measured the fractional CO_2_ concentration in incurrent air. From there, the air current was hydrated passing it through a bottle containing water (and several beads of Ascarite to keep CO_2_ levels low). The hydrated stream was then directed through egg-containing chambers in the MAVEn and back into the measurement side of the analyzer (referred to as cell B), which measured the fractional CO_2_ concentration in excurrent air. Data from the system was logged at 1 Hz using the MAVEn software.

Approximately every two weeks, each of the 16 eggs per species was weighed on an analytical balance (ME104TE/00, Mettler Toledo, Columbus, OH, USA) and subsequently placed individually in the 16 experimental chambers between the hours of 1600 and 2000 and left to run overnight until 0800-1000. Within each MAVEn cycle, air flow was first directed to the baseline channel for 5 min then to two experimental chambers sequentially for 20 min each and back to the baseline channel and so on through the remaining chambers.

### Analysis of metabolic data

Data were extracted and analyzed using R (v4.1.1, R Core Team, 2021). The raw data files contained the relative concentration of CO_2_ (parts per million, ppm) inside cell B compared with cell A (the reference cell) according to time (sampling frequency: 1 Hz). We converted raw measures (ppm) to molar rates of CO_2_ production (ṀCO_2_) using measured flow rates inside each chamber (which varied from 10 to 25 ml.min^−1^) and the Ideal Gas Law:

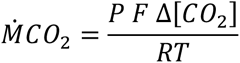

where ṀCO_2_ is the rate of CO_2_ production (mol.min^−1^), Δ[CO_2_] is the fractional concentration of CO_2_ in cell B relative to cell A, F is the flow rate (L.min^−1^), P is pressure (1 atm), R is the gas constant (0.08206 L.atm.K^−1^.mol^−1^) and T is the temperature measured by the MAVEn (K). Using custom scripts in R, we corrected the CO_2_ traces for baseline drift by subtracting the mean baseline values that bracket each set of two experimental chambers.

In general, egg metabolic rate increased continuously during development (Fig. 5A), which raises the questions of what representative values to use in other analyses. We approached this problem in two ways. First, for each egg, we estimated the metabolic rate halfway through the total developmental time from cubic regression splines and general additive models (GAM, *gam*: “mgcv”, k=5, bs= “cr”) fitted to the developmental trajectory of metabolism of each egg separately (Fig. S6). Second, for each egg, we estimated the total CO_2_ produced over its developmental period by integrating rates of CO_2_ production at each time point estimated by the GAM and multiplying that value by the total development time. A measure of mean metabolic rate was then obtained by dividing total CO_2_ produced by development time.

We tested the scaling relationships of mid-development metabolic rate and mean egg metabolic rate with egg mass by running linear mixed-effects regressions (*lmer*: “lme4”, ^67^ including species ID as a random effect. Variables were log_10_-transformed prior to analyses. 95% confidence intervals were computed to compare the estimated slope to isometry (slope *β*= 1). We then investigated how the total CO_2_ produced during embryogenesis (i.e., a proxy for the total energy for embryogenesis) varied with egg mass by running a similar linear mixed-effect model.

### Egg rates of water loss

To assess rates of water loss, we held freshly laid eggs of the five species for approximately two months in a Tupperware container at 75% relative humidity (at room temperature 21 – 23 °C). Humidity was controlled by filling the bottom of each container with saturated solutions of sodium chloride in water ^68^, and eggs were held in 24-well plates positioned above the salt solutions. For controls, we used the eggs held in 100% relative humidity on which we measured metabolic rates. Depending on their size, eggs were weighed twice per week on either a Mettler Toledo ME54TE/00 (± 0.1 mg) or a Sartorius MC-5 microbalance (± 1 µg). For each egg, we calculated water loss rate as the slope of the linear regression between egg mass and time. After two months, and before the eggs hatched, we transferred them to a drying oven (60 °C) for two weeks and reweighed them dry. From these data and the initial fresh masses, we calculated initial water contents. We tested the scaling relationships of egg water loss rate and initial egg water content with initial egg mass by performing a linear mixed-effects regression after log_10_- transformation, including species ID as a random effect. The effect of oviposition mode was added in the model with water loss as it appeared to have a large effect. For each species, we calculated an average rate of mass loss using a linear mixed-effects regression between egg mass and time, including egg ID as a random effect. At 100% relative humidity, in *E. calcarata* (slope *β*= -0.01 ± 0.01 mg.day^−1^, Wald *χ*^2^ test: *χ*^2^= 1.03, df = 1, p= 0.3) and in *M. extradentata* (*β*= -0.0008 ± 0.0007 mg.day^-1^, *χ*^2^= 1.54, df= 1, p= 0.21), egg mass did not significantly decrease over time. In contrast, for *H. dilatata* (*β*= -0.03 ± 0.002 mg.day^-1^, *χ*^2^= 176.8, df= 1, p< 0.0001), *P. philippinicum* (*β*= - 0.002 ± 0.0006 mg.day^−1^, *χ*^2^ = 9.38, df= 1, p= 0.002) and *E. tiaratum* (*β*= -0.005 ± 0.0007 mg.day^-1^, *χ*^2^= 43.9, df= 1, p< 0.0001), we found a significant decrease of mass over time. This decrease may be attributed to loss of organic matter through catabolism or to water loss during flow-through respirometry. However, mass loss rates estimated through the same method for eggs held at 75% relative humidity, showed much higher values (*H. dilatata*: *β*= -0.11 ± 0.002 mg.day^-1^, *P. philippinicum*: *β*= -0.008 ± 0.001 mg.day^-1^, *E. tiaratum*: *β*= -0.013 ± 0.001 mg.day^-1^), suggesting that most of the mass lost by eggs at 75% relative humidity were due to water loss.

## Results

### Phylogenetic analysis of transitions in oviposition style

Our ancestral state reconstruction unambiguously indicated that the ancestor lineages of Phasmatodea and Euphasmatodea dropped or flicked their eggs to the ground, like most extant phasmid species worldwide (Fig. 1). The associated transition matrix identified asymmetric transition rates between oviposition modes and suggested that transitions from dropping/flicking to burying (at least 16 times) or gluing (at least 7 times), and from gluing to burying (at least 3 times) were the most frequent. Only one reverse transition back to dropping/flicking from burying was recovered, although equivocally, in the New Zealand/New Caledonia clade (belonging to Lanceocercata, Fig. 1) highlighting that reverse transitions from more derived styles (i.e., burying or gluing) were very unlikely. Morphologically- and ecologically-diverse clades that colonized diverse habitats often exhibited multiple such transitions and diverse oviposition styles (e.g., Pseudophasmatinae, African/Malagasy clade, Necrosciinae, Lanceocercata) unlike morphologically- and ecologically- homogeneous clades (e.g., Heteropterygidae, Phylliidae)^33^.

**Figure 1:**
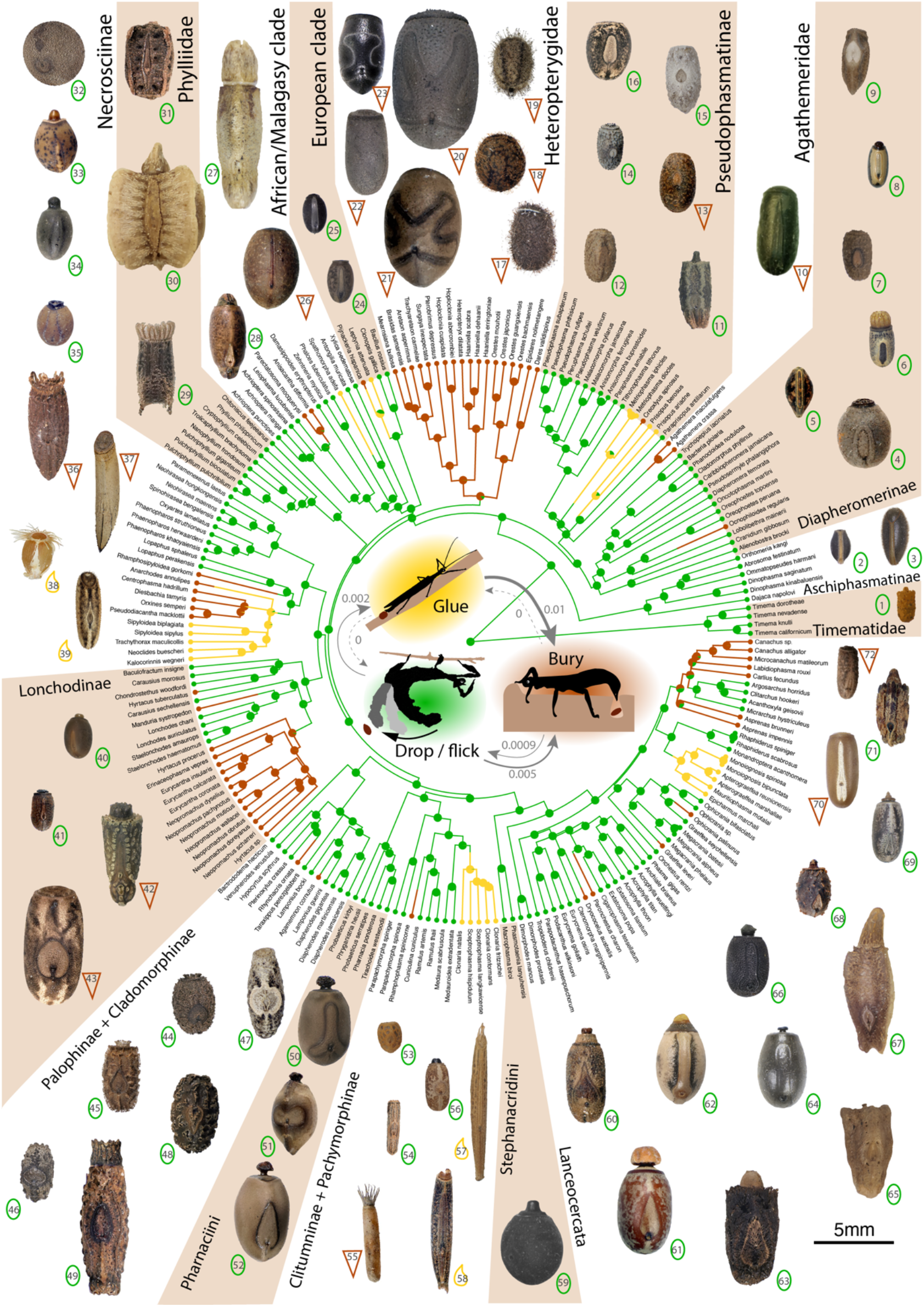
Egg morphological diversity and ancestral state reconstruction of oviposition mode in Phasmatodea. The ancestral state reconstruction used stochastic character mapping and a transition matrix (inset) estimated by maximum likelihood. Scaled egg pictures in dorsal view correspond to the species listed in table S1. Dropped or flicked eggs are represented by a green oval, buried eggs by a brown triangle and glued eggs by a yellow droplet.

### Size, life history, and pace of life

Across phasmids, larger females laid more and larger eggs (Table 1, Fig. 2A-B). After accounting for adult female body volume, we found a negative correlation (i.e., a trade-off) between lifetime egg number (fecundity) and egg volume (Table 1, Fig. 2C). Oviposition mode significantly affected the size-number tradeoffs. Females burying or gluing their eggs in specific locations, i.e., those that invest relatively more in parental care, appeared to lay fewer but relatively larger eggs relative to females that drop or flick their eggs away. Overall, lifetime reproductive output (lifetime fecundity × average egg volume) was strongly correlated with adult female volume but did not significantly differ between oviposition modes (Table 1, Fig. 2D). Larger eggs developed slower than smaller eggs and, for a given egg volume, glued eggs developed faster than buried or dropped eggs (Table 1, Fig. 2E).

**Figure 2:**
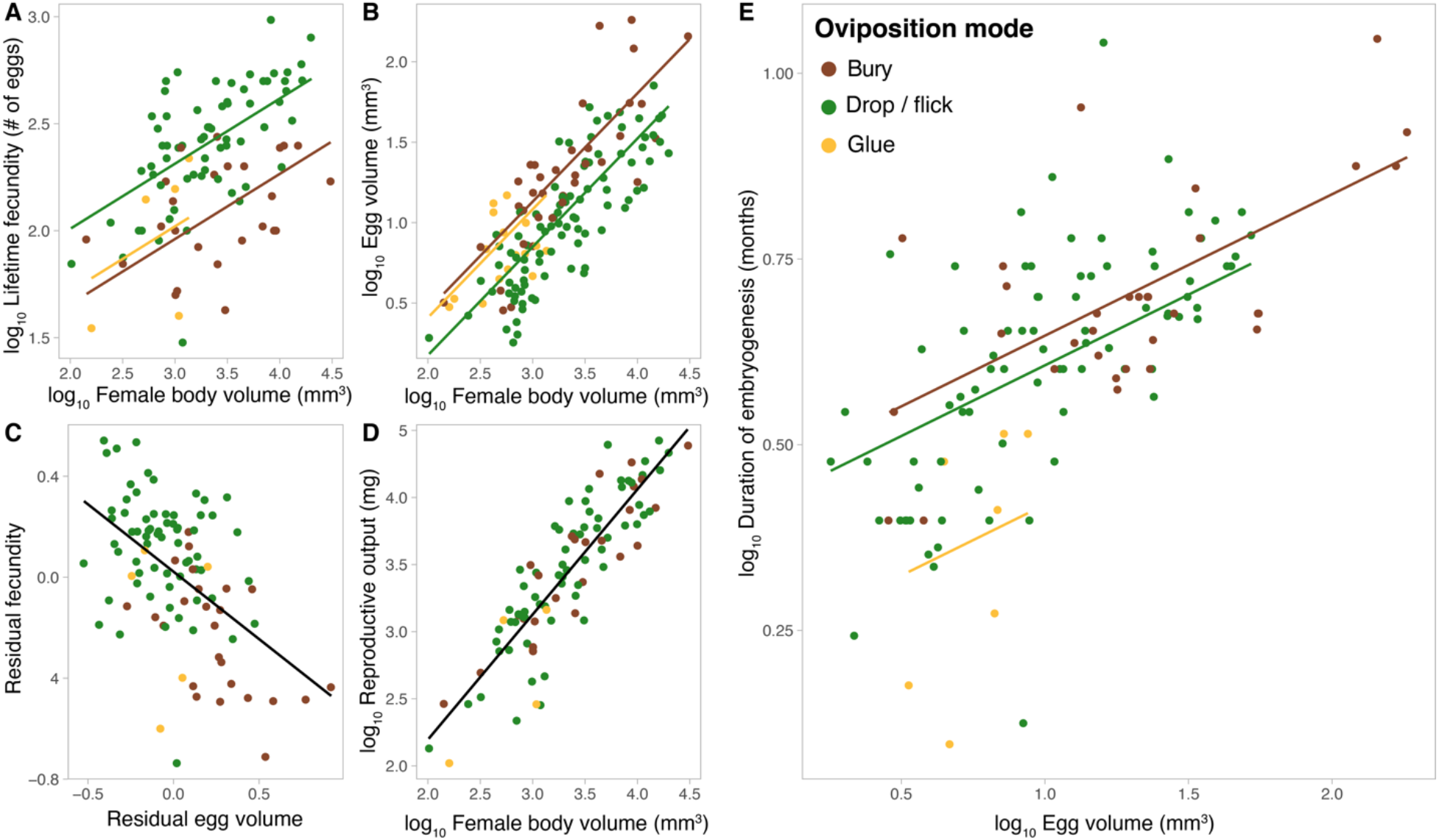
Life history predictors of egg size. Lifetime fecundity (**A**) and egg volume (**B**) compared to adult female body volume. **C**, Relationship between relative lifetime fecundity and relative egg volume after accounting for adult female body volume. **D**, Lifetime reproductive output (fecundity x egg volume) as a function of adult female body volume. **E**, Duration of embryogenesis, from egg laying to hatching, compared to egg volume. Colors represent different oviposition modes (see legend in **E**). Phylogenetic least square regressions are represented and correspond to the analyses reported in table 1.

**Table 1:** Life history correlates of egg size. The table presents the results of phylogenetic generalized least square models. The most likely value of Pagel’s lambda (phylogenetic signal) is presented along with type- I (sequential) ANOVA outputs and either estimated effect sizes or post-hoc pairwise comparisons between estimated marginal means using the Holm method to account for multiple testing, respectively for continuous or categorical explanatory variables. Significant effects are bolded (p<0.05).

| Response (n species) | Predictor | $\lambda$ | $F_{df1,df2}$ | P | Effect size ± Standard Error (continuous) or Post-hoc pairwise tests (Holm) (categorical) |
| --- | --- | --- | --- | --- | --- |
| $\log_{10}$ Lifetime fecundity (= egg number) (n = 96) | **$\log_{10}$ Adult female body volume** | 0 | $F_{1,92} = 44.27$ | **<.0001** | **$0.30 \pm 0.04$** |
| | Oviposition mode | | $F_{2,92} = 23.29$ | **<.0001** | **Drop – bury = $0.35 \pm 0.05$, $p < .0001$**<br>**Drop – glue = $0.29 \pm 0.11$, $p = 0.017$**<br>Bury – glue = $-0.06 \pm 0.12$, $p = 0.61$ |
| $\log_{10}$ Egg volume (n = 143) | **$\log_{10}$ Adult female body volume** | 0.57 | $F_{1,139} = 192.2$ | **<.0001** | **$0.67 \pm 0.05$** |
| | Oviposition mode | | $F_{2,139} = 15.3$ | **<.0001** | **Drop – bury = $-0.28 \pm 0.05$, $p < .0001$**<br>**Drop – glue = $-0.23 \pm 0.08$, $p = 0.005$**<br>Bury – glue = $0.05 \pm 0.08$, $p = 0.55$ |
| $\log_{10}$ lifetime fecundity (n = 96) | **$\log_{10}$ Adult female body volume** | 0.02 | $F_{1,93} = 40.1$ | **<.0001** | **$0.64 \pm 0.07$** |
| | **$\log_{10}$ Egg volume** | | $F_{1,93} = 33.9$ | **<.0001** | **$-0.53 \pm 0.09$** |
| $\log_{10}$ Reproductive output (egg number × volume) (n = 96) | **$\log_{10}$ Adult female body volume** | 0.12 | $F_{1,92} = 290.9$ | **<.0001** | **$0.91 \pm 0.06$** |
| | Oviposition mode | | $F_{2,92} = 2.56$ | 0.08 | Drop – bury = $0.04 \pm 0.06$, $p = 0.50$<br>Drop – glue = $0.28 \pm 0.13$, $p = 0.08$<br>Bury – glue = $0.24 \pm 0.14$, $p = 0.15$ |
| $\log_{10}$ Duration of embryogenesis (n = 121) | **$\log_{10}$ Egg volume** | 1 | $F_{1,117} = 45.3$ | **<.0001** | **$0.19 \pm 0.03$** |
| | Oviposition mode | | $F_{2,117} = 6.53$ | **0.002** | Drop – bury = $-0.04 \pm 0.04$, $p = 0.26$<br>**Drop – glue = $0.19 \pm 0.06$, $p = 0.008$**<br>**Bury – glue = $0.23 \pm 0.06$, $p = 0.001$** |

### Mechanical constraints on egg size and shape

A phylogenetic principal component analysis revealed that most of the variation in egg capsule shape pertained to variation in aspect ratio (i.e., relative elongation): PC1 (66% of the total variation) reflected variation in elongation, PC2 (21% of the total variation) reflected variation in axis of elongation (anterior-posterior or dorsal-ventral) (Fig. 3A, S1B-C). Most phasmid eggs were clustered in the center of the resulting egg morphospace, exhibiting a generic barrel-shaped egg capsule (Fig. 3A; e.g., *Anisomorpha buprestoides*, egg #12; *Diapherodes martinicensis*, egg #48; *Acrophylla wuelfingi*, egg #66 in Fig. 1). In contrast, other species had very elongated shapes (i.e., high PC1; e.g., *Sceptrophasma hispidulum*, egg #58 in Fig.1) or flattened lentil-shaped capsules (i.e., low PC2; e.g., *Oreophoetes peruana*, egg #5 in Fig. 1).

**Figure 3:**
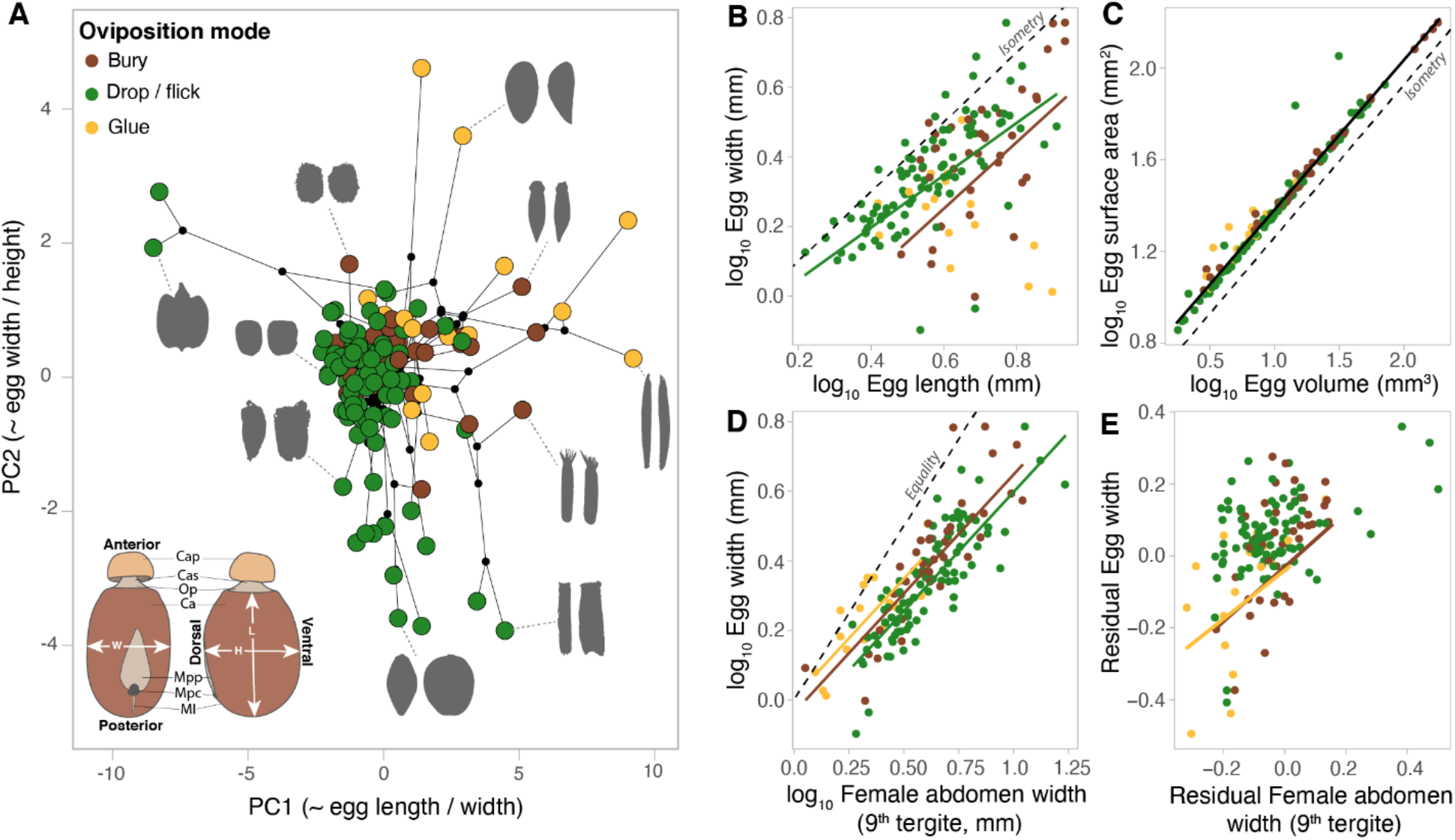
Egg shape and allometric scaling. **A**, Phasmid egg morphospace showing the two first axes of a phylogenetic principal component analysis. Black lines between points represent the underlying phylogenetic relationships between species. Black dots represent the inferred position of internal nodes. Egg silhouettes are represented in dorsal (left) and side (right) view. Bottom left inset shows the drawing of a phasmid egg (*Eurycnema osiris*, Lanceocercata): Cap, capitulum; Cas: capitulum stalk; Op, operculum; Ca, egg capsule; Mpp, micropylar playe; Mpc, micropylar cup; Ml, median line; L, egg length; W, egg width; H, egg height. **B**, Egg width as a function of egg length. **C**, Egg surface area as a function of egg volume. **D**, Egg width as a function of the width the adult female ninth tergite (i.e., where the egg is released). **E**, Residual egg width (i.e., residuals from the PGLS regression between egg width and egg length) compared to residual width of female ninth tergite (i.e., residuals from the PGLS regression between width of ninth tergite and adult female body volume). Phylogenetic least square regressions are represented in **B-E** and correspond to the analyses reported in table 1. Colors correspond to oviposition mode (see legend in **A**). Dashed lines represent isometric slopes (arbitrary intercept) in **B** and **C**, or the equality line in **D**.

Egg width was positively correlated with egg length in dropped and buried eggs but not in glued eggs (Table 2, Fig. 3B). The scaling relationship between egg width and egg length was significantly hypoallometric in dropped eggs (*β* = 0.78 ± 0.07, 95% confidence interval = [0.64; 0.92], isometric slope = 1) but did not differ from isometry in buried eggs (*β* = 1.03 ± 0.18, 95% confidence interval = [0.68; 1.39], isometric slope = 1). Buried eggs were relatively more elongated than dropped eggs (Table 2, Fig. 3B) while glued eggs ranged from the most elongated eggs (e.g., *Clonaria conformans,* egg #57 in Fig. 1) to more spherical eggs, notably in species where females glue their eggs in batches (Fig. 3A; e.g., *Trachythorax maculicollis,* egg #38 in Fig. 1). This morphological diversity suggests a strong influence of substrate properties on which eggs are glued (e.g., bark, grass, leaves) and laying strategy (e.g., eggs laid singly or in batches) on the shape of glued eggs.

**Table 2:** Allometric scaling of egg shape and mechanical constraints. The table presents the results of phylogenetic generalized least square models (PGLS). The most likely value of Pagel’s lambda (phylogenetic signal) is presented along with type-I (sequential) ANOVA outputs and either estimated effect sizes or post-hoc pairwise comparisons between estimated marginal means using the Holm method to account for multiple testing, respectively for continuous or categorical explanatory variables. Residual egg width corresponds to the residuals of a PGLS regression between egg width and egg length and therefore represents a measure of aspect ratio with lower values indicating more elongated eggs. Residual female ninth tergite width (i.e., the putative maximum diameter of the oviduct opening) corresponds to the residuals of a PGLS regression between ninth tergite width and female volume and therefore represents the relative elongation of the apex of the female’s abdomen, with lower values indicating a narrower abdomen. Significant effects are bolded (p<0.05).

| Response<br>(n species) | Predictor | $\lambda$ | $F_{df1,df2}$ | P | Effect size ± Standard Error (continuous)<br>OR<br>Post-hoc pairwise tests (Holm) (categorical) |
| --- | --- | --- | --- | --- | --- |
| log<sub>10</sub> Egg width<br>(n = 143) | log<sub>10</sub> Egg length | 0.97 | $F_{1,137} = 107.6$ | **<.0001** | **0.95 ± 0.16** |
| | Oviposition mode | | $F_{2,137} = 7.53$ | **0.0008** | Intercept pairwise comparisons:<br>Drop – bury = **0.10 ± 0.03, $p=0.005$**<br>Drop – glue = **0.11 ± 0.04, $p=0.005$**<br>Bury – glue = **0.02 ± 0.04, $p=0.63$** |
| | Interaction | | $F_{2,137} = 8.65$ | **0.0003** | Slope pairwise comparisons:<br>Drop – bury = **-0.20 ± 0.18, $p=0.25$**<br>Drop – glue = **0.87 ± 0.23, $p=0.0005$**<br>Bury – glue = **1.08 ± 0.27, $p=0.0003$** |
| log<sub>10</sub> Egg surface area<br>(n = 143) | log<sub>10</sub> Egg volume | 1 | $F_{1,137} = 3913$ | **<.0001** | **0.64 ± 0.02** |
| | Oviposition mode | | $F_{2,137} = 2.78$ | 0.07 | Intercept pairwise comparisons:<br>Drop – bury = **-0.02 ± 0.01, $p=0.16$**<br>Drop – glue = **-0.02 ± 0.02, $p=0.84$**<br>Bury – glue = **0.005 ± 0.02, $p=0.84$** |
| | Interaction | | $F_{2,137} = 1.08$ | 0.34 | Slope pairwise comparisons:<br>Drop – bury = **0.03 ± 0.02, $p=0.73$**<br>Drop – glue = **0.06 ± 0.05, $p=0.73$**<br>Bury – glue = **0.04 ± 0.05, $p=0.73$** |
| log<sub>10</sub> Egg width<br>(n = 142) | log<sub>10</sub> Female 9<sup>th</sup> tergite width | 0.57 | $F_{1,138} = 242.8$ | **<.0001** | **0.68 ± 0.04** |
| | Oviposition mode | | $F_{2,138} = 6.83$ | **0.0015** | Drop – bury = **-0.05 ± 0.02, $p=0.03$**<br>Drop – glue = **-0.09 ± 0.03, $p=0.004$**<br>Bury – glue = **-0.04 ± 0.03, $p=0.35$** |
| Residual egg width<br>(after accounting for egg length)<br>(n = 142) | Residual female 9<sup>th</sup> tergite width<br>(after accounting for female volume) | 0.95 | $F_{1,136} = 29.55$ | **<.0001** | **0.76 ± 0.19** |
| | Oviposition mode | | $F_{1,136} = 7.22$ | **0.001** | Intercept pairwise comparisons:<br>Drop – bury = **0.07 ± 0.02, $p=0.01$**<br>Drop – glue = **0.07 ± 0.04, $p=0.07$**<br>Bury – glue = **0.005 ± 0.04, $p=0.88$** |
| | Interaction | | $F_{1,136} = 4.41$ | **0.014** | Slope pairwise comparisons:<br>Drop – bury = **-0.51 ± 0.21, $p=0.044$**<br>Drop – glue = **-0.43 ± 0.19, $p=0.046$**<br>Bury – glue = **0.08 ± 0.24, $p=0.73$** |

Overall, egg surface area scaled isometrically with egg volume (*β* = 0.65 ± 0.01, 95% confidence interval = [0.633; 0.674], isometric slope = 0.667) and this relationship was not affected by oviposition mode (Table 2, Fig. 3C).

Egg width scaled hypoallometrically with the width of the adult female’s ninth tergite, where the opening of the oviduct is located (i.e., the putative maximum egg width for the egg to be able to come out of the oviduct) (Table 2, Fig. 3D; *β* = 0.68 ± 0.04, 95% confidence interval = [0.60; 0.77], isometric slope = 1). The width of almost all eggs was smaller than that of the ninth tergite (Fig. 3D). Glued and buried eggs were relatively wider and their width was closer to the width of the female ninth tergite, sometimes slightly larger suggesting potential dilatation of the oviduct opening (Table 2, Fig. 3D). Residual egg width after accounting for egg length (i.e., reflecting aspect ratio) was positively correlated with residual female ninth tergite width, after accounting for female volume, in species that bury or glue their eggs but not significantly in species that drop or flick them away (Table 2, Fig. 3E). Thus, females with a relatively narrower abdominal extremity (i.e., with a relatively more elongated body) lay relatively more elongated eggs but this relationship was only true for species that bury or glue their eggs, which tend to be relatively larger (Fig. 2B) and closer to the limit in egg width imposed by the oviduct opening diameter (Fig. 3D). This suggests that, in species with more derived oviposition modes and relatively larger eggs, the width of these eggs is mechanically constrained by the diameter of the female oviduct opening.

### Response to ecological circumstances

After accounting for the effects of adult female volume, lifetime fecundity and oviposition mode, we found significant effects of female flight capacity and climate PC1 on egg volume (Table 3). Flight capable females and females living in more temperate and seasonal regions (reflected by climate PC1) laid relatively smaller eggs (Table 3).

**Table 3:** Effect of ecological variables on egg size and shape. The table presents the results of phylogenetic generalized least square models (PGLS). The most likely value of Pagel’s lambda (phylogenetic signal) is presented along with type-I (sequential) ANOVA outputs and either estimated effect sizes or post-hoc pairwise comparisons between estimated marginal means using the Holm method to account for multiple testing, respectively for continuous or categorical explanatory variables. Significant effects are bolded (p<0.05).

| Response (n species) | Predictor | $\lambda$ | $F_{df1,df2}$ | P | Effect size ± Standard Error (continuous) or Post-hoc pairwise tests (Holm) (categorical) |
| --- | --- | --- | --- | --- | --- |
| log<sub>10</sub> Egg volume (n = 96) | log<sub>10</sub> Female body volume | 0.44 | **$F_{1,88} = 182.6$** | **<.0001** | **$0.66 \pm 0.05$** |
|  | log<sub>10</sub> lifetime fecundity |  | **$F_{1,88} = 34.1$** | **<.0001** | **$-0.32 \pm 0.09$** |
| | Oviposition mode | | **$F_{2,88} = 6.47$** | **0.002** | Drop – bury = $-0.14 \pm 0.06$, $p=0.09$<br>Drop – glue = $-0.07 \pm 0.11$, $p=1$<br>Bury – glue = $0.07 \pm 0.11$, $p=1$ |
|  | Adult female flight capacity |  | **$F_{1,88} = 7.27$** | **0.008** | **Flying – flightless = $-0.21 \pm 0.08$, $p=0.007$** |
|  | Climate PC1<br>(~Net primary production, annual temperature and precipitation) |  | **$F_{1,88} = 10.2$** | **0.002** | **$0.04 \pm 0.01$** |
| | Climate PC2<br>(~Precipitation seasonality) | | $F_{1,88} = 2.45$ | 0.12 | $-0.03 \pm 0.02$ |
| log<sub>10</sub> Egg surface area (n = 96) | log<sub>10</sub> Egg volume | 0.72 | **$F_{1,88} = 1801.0$** | **<.0001** | **$0.65 \pm 0.02$** |
| | log<sub>10</sub> lifetime fecundity | | $F_{1,88} = 0.03$ | 0.86 | $0.006 \pm 0.02$ |
| | Oviposition mode | | $F_{2,88} = 0.31$ | 0.73 | Drop – bury = $-0.009 \pm 0.02$, $p=1$<br>Drop – glue = $-0.015 \pm 0.03$, $p=1$<br>Bury – glue = $-0.006 \pm 0.11$, $p=1$ |
| | Adult female flight capacity | | $F_{1,88} = 0.04$ | 0.84 | **Flying – flightless = $0.004 \pm 0.02$, $p=0.86$** |
| | Climate PC1<br>(~Net primary production, annual temperature and precipitation) | | $F_{1,88} = 0.79$ | 0.38 | $0.003 \pm 0.003$ |
| | Climate PC2<br>(~Precipitation seasonality) | | $F_{1,88} = 1.16$ | 0.28 | $-0.005 \pm 0.005$ |
| log<sub>10</sub> Egg width<br>(n = 143) | log<sub>10</sub> Egg length | 0.98 | F<sub>1,136</sub> = 104.7 | <.0001 | 0.65 ± 0.07 |
|  | Oviposition mode |  | F<sub>2,136</sub> = 6.94 | 0.001 | Drop – bury = 0.08 ± 0.03, p=0.009<br>Drop – glue = 0.11 ± 0.04, p=0.009<br>Bury – glue = 0.03 ± 0.04, p=0.39 |
|  | Adult female flight capacity |  | F<sub>1,136</sub> = 1.39 | 0.24 | Flying – flightless = -0.03 ± 0.03, p=0.23 |
|  | Climate PC1<br>(~Net primary production, annual temperature and precipitation) |  | F<sub>1,136</sub> = 7.54 | 0.007 | 0.01 ± 0.005 |
|  | Climate PC2<br>(~Precipitation seasonality) |  | F<sub>1,136</sub> = 2.76 | 0.10 | -0.01 ± 0.007 |
| log<sub>10</sub> Reproductive output<br>(egg number × volume)<br>(n = 96) | log<sub>10</sub> Female body volume | 0.13 | F<sub>1,89</sub> = 319.4 | <.0001 | 0.86 ± 0.06 |
|  | Oviposition mode |  | F<sub>2,89</sub> = 2.85 | 0.06 | Drop – bury = 0.096 ± 0.065, p=0.43<br>Drop – glue = 0.15 ± 0.14, p=0.57<br>Bury – glue = 0.05 ± 0.14, p=0.72 |
|  | Adult female flight capacity |  | F<sub>1,89</sub> = 4.40 | 0.04 | Flying – flightless = -0.22 ± 0.10, p=0.025 |
|  | Climate PC1<br>(~Net primary production, annual temperature and precipitation) |  | F<sub>1,89</sub> = 8.64 | 0.004 | 0.04 ± 0.01 |
|  | Climate PC2<br>(~Precipitation seasonality) |  | F<sub>1,89</sub> = 0.20 | 0.66 | -0.01 ± 0.02 |
| log<sub>10</sub> Duration of embryogenesis<br>(under 20-24°C)<br>(n = 121) | log<sub>10</sub> Egg volume | 1 | F<sub>1,114</sub> = 44.2 | <.0001 | 0.19 ± 0.03 |
|  | Oviposition mode |  | F<sub>2,114</sub> = 6.38 | 0.002 | Drop – bury = -0.04 ± 0.04, p=0.28<br>Drop – glue = 0.19 ± 0.07, p=0.009<br>Bury – glue = 0.23 ± 0.07, p=0.002 |
|  | Adult female flight capacity |  | F<sub>1,114</sub> = 0.08 | 0.77 | Flying – flightless = 0.009 ± 0.03, p=0.75 |
|  | Climate PC1<br>(~Net primary production, annual temperature and precipitation) |  | F<sub>1,114</sub> = 0.01 | 0.91 | 0.0009 ± 0.006 |
|  | Climate PC2<br>(~Precipitation seasonality) |  | F<sub>1,114</sub> = 0.15 | 0.70 | -0.003 ± 0.007 |

After accounting for egg volume, egg surface area was not significantly affected by any of our ecological predictors. Therefore, species experiencing drier conditions, either seasonally (i.e., high climate PC2) or constantly (i.e., low climate PC1) do not seem to minimize egg surface to volume ratio to limit water loss.

After accounting for egg length, relative egg width (i.e., reflecting aspect ratio) was significantly affected by oviposition mode and climate PC1 (Table 3). Glued and buried eggs were more elongated than dropped eggs and eggs of species found in drier and more temperate and seasonal regions were slightly rounder (Table 3).

Total female reproductive output was significantly affected by female flight capacity and climate PC1, even after accounting for female body volume (Table 3). Flightless females were able to invest relatively more in egg production than were flight capable ones. This suggests either a physiological cost of the flight apparatus or a cost of carrying more and/or larger eggs for flying, and potentially explains why even after accounting for relative fecundity, flightless females lay relatively larger eggs. Similarly, females in regions with a higher net primary productivity (and likely benefitting from greater food availability) were able to invest relatively more in egg production, also suggesting that females in temperate and less productive regions may be resource-limited compared to those from tropical regions. This could also potentially explain why tropical females lay relatively larger eggs even after accounting for relative fecundity. However, we note that our fecundity data were mostly obtained from breeding cultures, in which insects are fed *ad libitum*. Alternatively, differences in total reproductive investment may reflect reduced adult life spans in seasonal regions with short periods of favorable conditions.

Finally, after accounting for variation in egg volume, the duration of embryogenesis was affected only by oviposition mode, with glued eggs developing relatively faster than buried and dropped eggs (Tables 1, 3). The other ecological variables had no significant effects (Table 3).

However, our data on the duration of egg development were obtained from breeding cultures in which eggs are usually incubated in stable room temperatures (20-24°C) and high humidities (>80%). Therefore, the effects of climate variables may be masked, especially if duration of embryogenesis is highly sensitive to temperature. Nevertheless, this outcome suggests that adaptation to local macroclimatic conditions may not significantly affect the evolution of egg chorion properties (e.g., conductance to water and respiratory gases) or other egg physiological properties. Indeed, such genetic adaptations should lead to differences in egg development duration among species adapted to contrasting climatic conditions once they are held in common conditions in the laboratory. In contrast, microclimatic conditions are likely to play a much larger role. Despite being held in very similar conditions, eggs from species that oviposit onto or into the soil developed more slowly than those glued onto substrates, that likely experience very different microclimatic conditions in the wild.

### Phylogenetic path analyses

Phylogenetic path analyses were used to infer the most likely causal relationships among egg size and shape, and variables relating to life history, mechanical constraints and ecology and behavior (Fig. 4). This analysis revealed that variation in egg size has been directly driven by many variables related to all three hypothesis categories: tropical climates, larger body size, relatively wider abdomens, and flight loss in females favored the evolution of larger eggs. Inversely, variation in egg size has also been causing variation in various life history and behavioral traits: larger eggs reduce female fecundity (total egg number), lengthen embryonic development, and drive the evolution of parental care (i.e., oviposition in specific locations). In parallel, egg shape also has evolved in response to changes in several other traits: smaller and thinner females produce more elongated eggs, likely highlighting mechanical constraints for the egg to make its way through the female reproductive canal. Females burying or gluing their eggs produce more elongated eggs, likely facilitating their insertion into substrates through their streamlined shape, or adhesion to surfaces through increased contact area.

**Figure 4:**
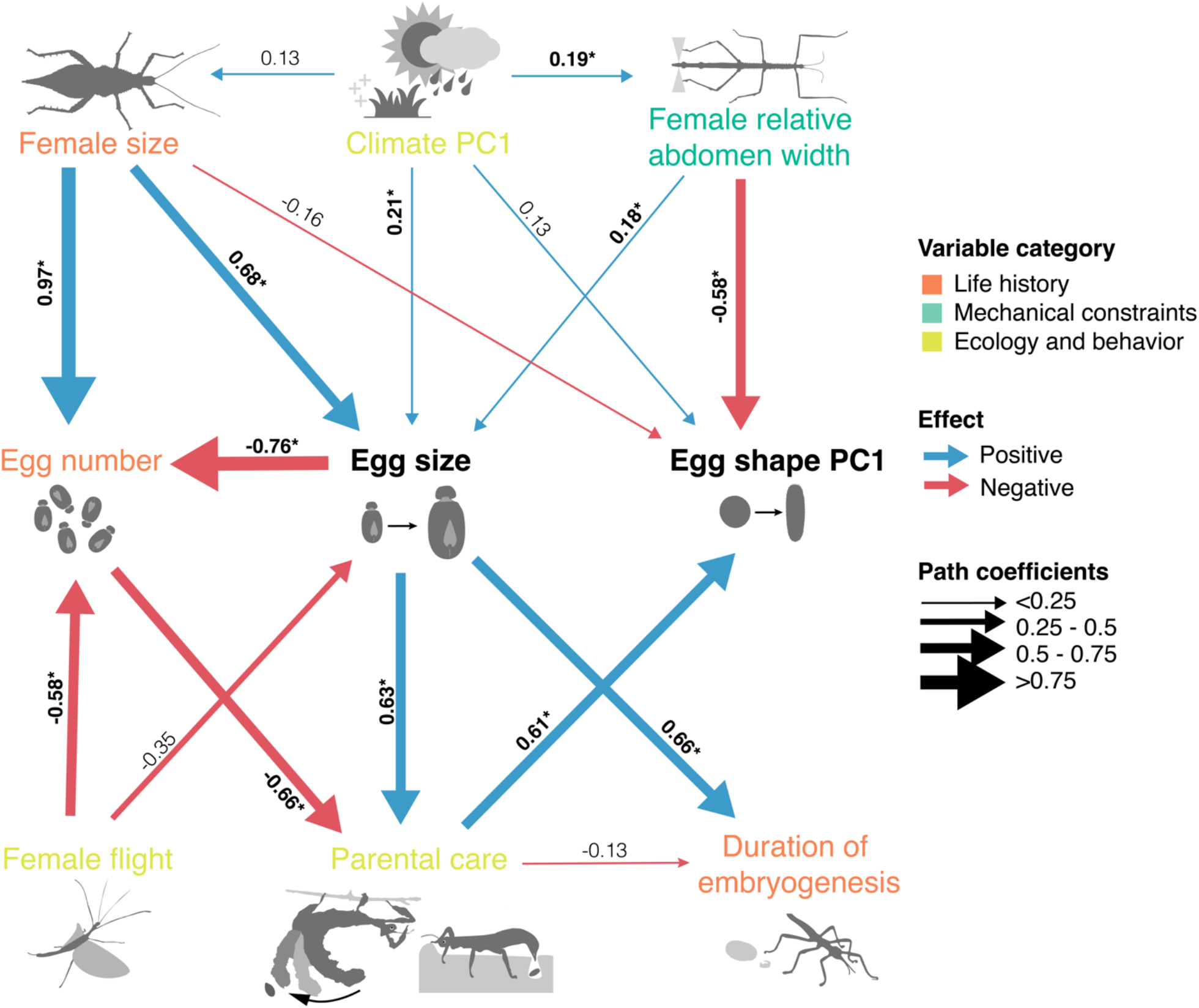
Averaged best-fitting path models (ΔCICc <2). Arrows show the direction of the path. Their color and width represent direction and size of regression coefficient (adjacent to arrows). Coefficients significantly deviating from 0 are written in bold with an asterisk.

### Egg metabolic rate, energy use and water loss

The five sampled species differed extensively in egg size and consequently in duration of embryogenesis, from 3.7 mg (average 4.07 ± 0.07 mg) and 94 days (average 98±0.9 days) in *Medauroidea extradentata*, to 173 mg (average 155.9±2.4 mg) and 377 days (average 341±6.8 days) in *Heteropteryx dilatata*. Egg metabolic rate increased exponentially during embryonic development in all five species (Fig. 5A). Mid-development and mean metabolic rate scaled hypoallometrically with egg mass across species (respectively *β* = 0.70 ± 0.16, 95% confidence interval: [0.36 ; 0.98], and *β* = 0.78 ± 0.10, [0.58 ; 0.97]; Table 4, Fig. 5B), indicating that larger eggs have a relatively lower metabolic rate than smaller eggs. Estimated cumulative CO_2_ produced during embryogenesis scaled isometrically with egg mass (Table 4, Fig. 5C), suggesting that larger eggs use proportionately more energy than smaller eggs for development. In all five species, eggs lost water at a constant rate during embryogenesis (Fig. 5D), and this rate increased hypoallometrically with egg mass after accounting for oviposition mode (Table 4, Fig. 5E). The recovered scaling exponent (0.29 ± 0.07, [0.15 ; 0.43]) was significantly lower than the expected scaling exponent of egg surface area with egg mass (0.66) suggesting that larger eggs have reduced chorion permeability, possibly from increased thickness or changes in layer properties ^69^. Interestingly, water loss rate was relatively higher in the two largest species, which bury their eggs. The three others drop their eggs (Table 4, Fig. 5E). While the effect of size and oviposition mode may be confounded, these patterns suggest that buried eggs are relatively more permeable to water. Finally, we found a strong isometric relationship between egg water content and egg mass (Table 4, Fig. 5F), suggesting that small and large eggs contain the same proportion water (49 ± 0.01%).

**Table 4:** Allometry of egg energy use and water loss. The table presents the results of linear mixed-effects models accounting for species as a random effect. Effect sizes (i.e., scaling exponents) are indicated along with type II ANOVA (Wald *χ*^2^) outputs including p values.

| Response | Fixed effect | Random effect | Effect size ± Standard Error<br>[95% Confidence interval] | Wald $\chi^2_{df}$ | P |
| --- | --- | --- | --- | --- | --- |
| $\log_{10}$ Mid embryonic development metabolic rate | $\log_{10}$ Egg mass | species | $0.70 \pm 0.16$<br>[0.36 ; 0.98] | $\chi^2_1 = 19.9$ | <.0001 |
| $\log_{10}$ Mean embryonic development metabolic rate | $\log_{10}$ Egg mass | species | $0.78 \pm 0.10$<br>[0.58 ; 0.97] | $\chi^2_1 = 62.9$ | <.0001 |
| $\log_{10}$ Total CO<sub>2</sub> produced during embryogenesis | $\log_{10}$ Egg mass | species | $0.97 \pm 0.09$<br>[0.76 ; 1.16] | $\chi^2_1 = 108.3$ | <.0001 |
| $\log_{10}$ Water loss rate | $\log_{10}$ Egg mass | species | $0.29 \pm 0.07$<br>[0.15 ; 0.43] | $\chi^2_1 = 16.0$ | <.0001 |
| | Oviposition mode | | Drop – Bury : $-0.81 \pm 0.08$<br>[-0.97 ; -0.65] | $\chi^2_1 = 95.7$ | <.0001 |
| $\log_{10}$ Water content | $\log_{10}$ Egg mass | species | $0.99 \pm 0.02$<br>[0.96 ; 1.03] | $\chi^2_1 = 2867$ | <.0001 |

**Figure 5:**
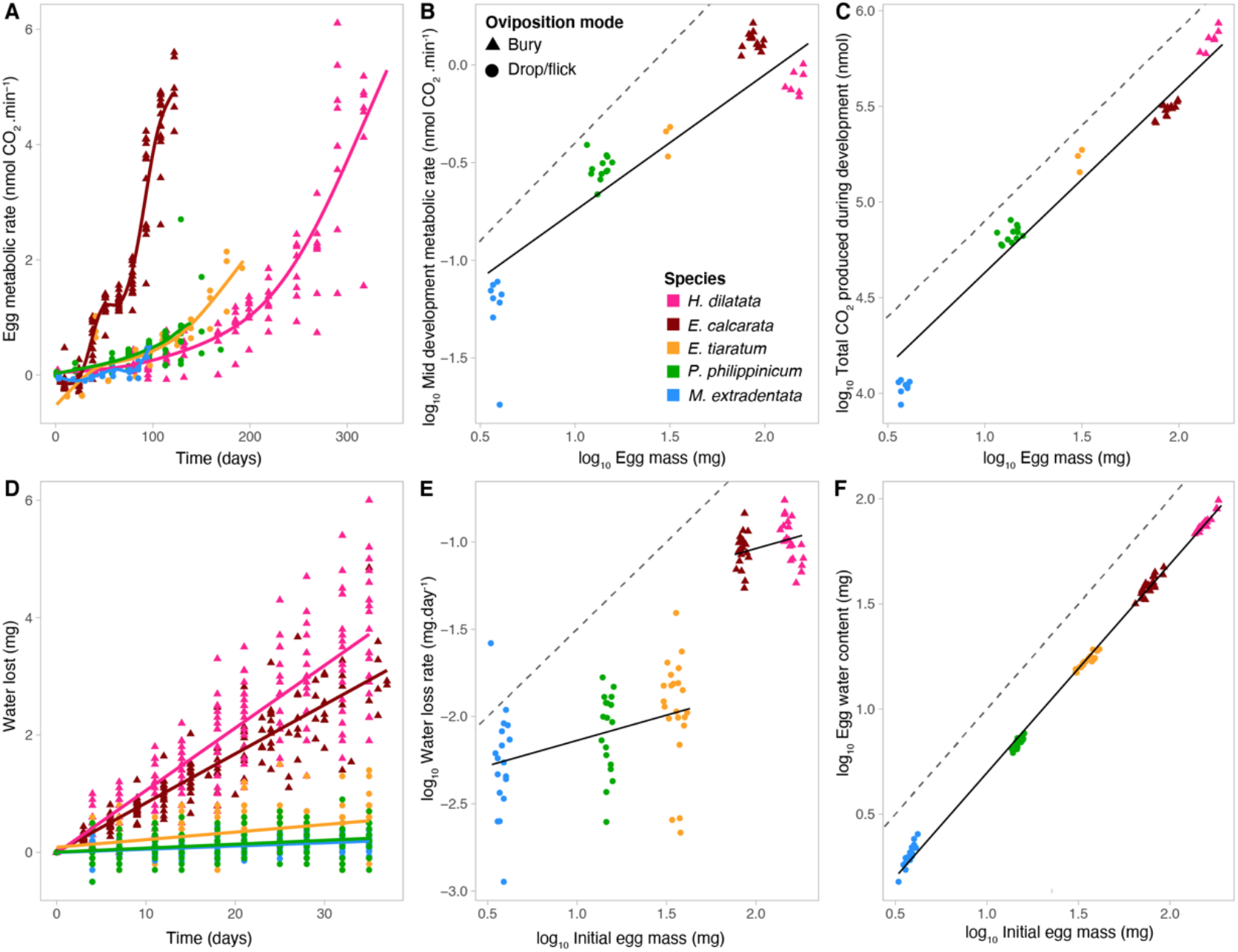
Allometry of egg energy use and water loss. **A.** Egg metabolic rate as a function of time (until hatching). Lines depict smoothing generalized additive models for each species. **B.** Estimated mid embryogenesis metabolic rate compared to egg mass. **C.** Estimated total CO2 produced during embryogenesis as a function of egg mass. **D.** Cumulative egg water loss as a function of time (until hatching). Lines depict linear regressions for each species. **E.** Water loss rate compared to initial egg mass (right after being laid). **F.** Initial egg water content as function of initial egg mass. Colors represent the five different species and shapes represent oviposition modes (see legend in **B**). Dashed lines represent isometry (arbitrary intercept). Solid black lines represent linear mixed-effects regressions (see table 4 for details on the statistical analyses).

## Discussion

Using a single mesodiverse clade of insects – the stick and leaf insects (order Phasmatodea, comprising approximately 3,400 described species) – we assessed a set of hypothesized drivers of egg diversification by combining phylogenetic analyses with physiological measurements of rates of metabolism and water loss. Together, the data suggest that egg morphological diversification is driven by a complex suite of factors including female resource allocation strategies, constraints arising from eggs interacting mechanically with various functions of the female body, the effects of female oviposition and locomotor (flight) behavior, and, to a lesser extent, the influences of climatic context in which lineages have evolved.

In our analyses, variation in egg size appeared mainly driven by female resource allocation strategies and trade-offs. Larger females lay more and larger eggs, but egg size strongly limits egg number (Fig. 2A-C, 4), providing strong evidence of egg size-number tradeoffs, consistent with previous studies on insects ^23,24^. The total reproductive investment of females (cumulative mass of laid eggs during their lifetime) appeared tightly related to female size (Fig. 2D). This highlights that females have access to limited resources for reproduction, which they may allocate among larger but fewer or smaller but more numerous eggs. Interestingly, we found a relatively strong negative effect of flight capacity on egg size and number (Fig. 4): flightless females had a higher lifetime reproductive investment as they laid relatively more and larger eggs compared to flight capable ones. The relatively reduced reproductive investment of flight capable females suggests that the physiological cost of wings and associated flight muscles may trade-off with reproductive output^63^. Alternatively, large eggs may impair the flight performance of females by increasing wing loading^70^. Stick and leaf insects are ideally suited to address how flight affects resource allocation strategies and egg diversification as they have undergone numerous shifts between winged forms and wingless forms ^48,60^. In accordance with Church, Donoughe et al.^8^, who also focused on Phasmatodea for this analysis, we did not find any effect of flight capacity on egg shape, in contrast with birds^3^.

Larger eggs also consistently required longer periods of time to develop. This result stands in striking contrast to the data reported by Church, Donoughe et al.^8^, who found no relationship between egg size and developmental duration. The difference likely reflects the phylogenetic scope of analysis ^71^. In their analysis, Church, Donoughe et al.^8^ used data on 66 genera distributed across seven major insect orders (both hemi- and holometabolous), such that order-specific differences may have obscured direct effects of egg size, if any, on duration of development. By contrast, our study, which analyzed 120 species distributed within a single order, finds strong evidence that larger eggs require longer developmental periods. This result is consistent with other reports on both insects ^9,12,13^ and vertebrates ^14–16^. Consistent with our observation that large eggs require longer developmental periods, egg size also had systematic effects on rates of egg metabolism and water loss. Specifically, both rates scaled hypoallometrically, such that large eggs had disproportionately low rates. Although the confidence intervals are wide due to the low number of species used (five), the values of the metabolic scaling coefficient (*β* = 0.70, 95% confidence interval: [0.36 ; 0.98] for mid-development metabolic rate; *β* = 0.78 [0.58 ; 0.97] for mean metabolic rate) lie in the range expected from a wide variety of other metabolic scaling studies (insect eggs: *β* = 0.92 [0.86; 0.98] ^19^, non-avian reptile eggs: *β* = 0.82 [0.77; 0.87] ^72^, avian eggs: *β* = 0.714 ± 0.09 (SE) ^73,74^, ectotherm eggs: *β* = 0.66 [0.65; 0.80] ^75^). The emerging picture is of larger eggs generally having longer developmental durations supported by lower mass-specific metabolic rates. There is scope, however, for additional covariation in metabolic rates and developmental duration beyond just the effects of size. For example, the two largest eggs in our experiment, *E. calcarata* (84.2 ± 0.8 mg) and *H. dilatata* (155.9 ± 2.4 mg), had relatively similar metabolic rates at the temporal midpoint of development (Fig. 5B). However, *E. calcarata* eggs develop much faster (121 ± 2 days versus 335 ± 5 days for *H. dilatata*), which is supported by a much more rapid increase in metabolism during the course of development (Figure 4A). Finally, the total energy devoted to development, as measured by estimated total CO_2_ emission over the entirety of development, scaled isometrically with egg size (Fig. 5C). Collectively, these observations suggest that egg size, developmental duration, rates of energy (and water) expenditure, and total energy reserves are all tightly correlated^76^. This outcome likely reflects the division of eggs into finite packets of energy and materials that exchange only water, carbon dioxide, and oxygen with their surroundings.

A major consequence of variation in relative egg size and number was the oviposition strategy adopted by females. Female phasmids either drop or flick their eggs away (the ancestral state; Fig. 1) or place them in specific locations, typically by gluing or burying them (two derived states that each have evolved at least five times within the clade). Larger and fewer eggs appeared to favor placement of the eggs in specific locations (Fig. 2A-C, 4), implying greater parental investment per offspring associated with the greater time investment and predator exposure involved in placing the eggs. Similar relationships between parental care and egg size have also been found in other ectotherms including fishes ^77^ and frogs ^78^. However, a recent study failed to find a relationship between egg size and the extent of parental care across 287 species of insects from 16 orders, and only found an effect on egg number ^79^. In addition, glued (but not buried) eggs evolved significantly shorter developmental durations for their size (Fig. 2E). Glued eggs present the advantage that offspring will hatch directly on the host plant, saving the time and effort associated with finding a suitable host plant and climbing it (e.g., *Spinosipyloidea doddi* females attach their hairy eggs on the hairy leaves of their unique and very tall host tree ^45^). However, they may be more vulnerable to egg predators and parasitoids, which, we hypothesize, could select for shorter development times ^80^.

Gluing and burying favored the evolution of more elongated eggs (Fig. 4). This effect may reflect one or more of several pressures: ease of oviposition – elongated eggs may be easier to bury within soil; camouflage – elongated eggs laid on leaves may be more difficult for visual predators to spot if they look less like the typically round eggs of most insect species or more like plant seeds ^27^; enhanced oxygen access – oxygen can be less concentrated in wet soils and elongated eggs may maximize their uptake by minimizing diffusion distances between surface and central tissues ^25,81^; contact surface area – glued elongated eggs may attach more strongly to substrates thanks to a larger contact area ^82,83^; and passage through the oviduct – elongated eggs may pass through the female oviduct more easily. Phasmids repeatedly evolved very elongated female body shapes, likely as a result of twig mimicry (masquerade)^33^. Consequently, the resulting narrow abdominal shapes appear to have favored smaller and more elongated eggs (Fig. 4), suggesting mechanical constraints imposed by these unusual body shapes. Females of species that glue or bury their eggs produce relatively larger eggs and therefore may be particularly affected by these constraints as the diameters of these eggs can approach or even exceed the diameters of the abdominal segments through which they pass during oviposition (Fig. 3D-E). Similarly, egg shape appears largely determined by anatomical constraints imposed by oviduct and pelvis width in birds^3,84^.

One surprising outcome of our analyses was the minimal predictive effects of climate variables. Phasmids are distributed across a broad range of tropical and temperate habitats ^85–87^, with significant variation in mean conditions and patterns of variation in major climate variables such as temperature, humidity, and rainfall (Fig. S3). A priori, one might expect that eggs in habitats with low annual precipitation would be larger and rounder to maximize water retention through a reduced surface to volume ratio; that eggs in more seasonal and potentially unfavorable environments would be larger to favor offspring survival and performance^88^; or that eggs in less productive environments (i.e., with a lower resource availability) would be smaller because of a reduced reproductive investment^89^. Indeed, we found evidence that females from temperate and drier areas associated with lower net primary productivity tended to be smaller and to lay slightly smaller and rounder eggs, even after accounting for female size and fecundity (Fig. 4). Thus, we show that these females invest relatively less in egg production (reproductive output), compared to females of tropical species. Overall, however, the effect sizes of the climate variables (collapsed into principal component axes) were small compared to the effect sizes of other predictors examined in this study (Table 3, Fig. 4). This outcome likely reflects that egg experience is poorly captured by the large-scale gridded climate data that we used (bin size 4.6 km on a side). Eggs experience microclimates rather than macroclimates, and conditions in or on the surface of the soil may differ strongly from conditions characterizing that grid cell ^90^. Likewise, eggs that are glued onto leaves experience conditions at least partly determined by leaf physiology ^91^.

Overall, our results on Phasmatodea align with the broad findings of Church, Donoughe et al. ^8^, notably in identifying a strong association between oviposition ecology and egg size and shape. In their study, ‘oviposition ecology’ referred primarily to whether insects laid their eggs in aerial environments or in aquatic environments, which characterize both insects that lay eggs in freshwater or parasitoids that lay eggs in the tissues of other animals. They found that eggs laid in aquatic environments were systematically smaller and more elongated. We were unable to test the consequences of evolutionary transitions between aerial and aquatic oviposition sites, as no phasmids oviposit in water or are parasitoids. Our data on phasmids further suggested that oviposition mode was a consequence of variation in egg size and that, in turn, it directly affected the evolution of egg shape. These results suggest that renewed attention should be directed toward understanding how ovipositing females experience interactions between eggs and their substrates, and the physical and biotic experiences that eggs have in their microsites.

## Conclusion

In conclusion, the staggering diversity of insect eggs is obvious, but we have lacked clear understanding of the major processes that underpin that diversification. Recent studies that leverage larger data sets, including this one, are starting to reveal drivers and consequences of this diversity. In particular, variation in egg size appears to be driven mainly by variation in female resource allocation strategy and trade-offs while egg shape appears to be influenced primarily by mechanical constraints imposed by the female body morphology and the oviposition mode and substrate.

## Acknowledgements

We thank François Tetaert for allowing us to use his image database of stick insect eggs (phasmatodea.fr); Garret Jolma for help in the lab developing early water-loss protocols; Brenna Shea and The Missoula Insectarium for providing fresh eggs of several phasmid species; Camille Thomas-Bulle and Douglas J. Emlen for insightful discussions on the manuscript.

